# Host Competitive Asymmetries Accelerate Viral Evolution in a Microbe-Virus Coevolutionary System

**DOI:** 10.1101/2025.03.09.642151

**Authors:** Armun Liaghat, Martin Guillemet, Rachel Whitaker, Sylvain Gandon, Mercedes Pascual

**Affiliations:** Department of Ecology and Evolution, University of Chicago, Chicago, IL, USA; Department of Biology, New York University, New York, NY, USA; Departmentof Environmental Science, New York University, New York, NY, USA; Institute for Integrative Biology, ETH Zürich, 8005 Zürich, Switzerland; CEFE, CNRS, Universite de Montpellier, EPHE, IRD, Montpellier, France; Carl R. Woese Institute for Genomic Biology, University of Illinois at Urbana-Champaign, Urbana, IL, USA; Department of Microbiology, University of Illinois at Urbana-Champaign, Urbana, IL, USA; Santa Fe Institute, Santa Fe, NM, USA

## Abstract

Microbial host populations evolve traits conferring specific resistance to viral predators via various defense mechanisms, while viruses reciprocally evolve traits to evade these defenses. Such co-evolutionary dynamics often involve diversification promoted by negative frequency-dependent selection. However, microbial traits conferring competitive asymmetries can induce directional selection, opposing diversification. Despite extensive research on microbe-virus co-evolution, the combined effect of both host trait types and associated selection remains unclear. Using a CRISPR-mediated co-evolutionary system, we examine how the co-occurrence of both trait types impacts viral evolution and persistence, previously shown to be transient and non-stationary in computational models. A stochastic model incorporating host competitive asymmetries via variation of intrinsic growth rates reveals that competitively-advantaged host clades generate the majority of immune diversity. Greater asymmetries extend viral extinction times, accelerate viral adaptation locally in time, and augment long-term local adaptation. These findings align with previous experiments, and provide further insights into long-term co-evolutionary dynamics.

## Introduction

Pangenomic analyses are increasingly revealing the breadth of both inter- and intra-specific diversity that co-occur in natural microbial populations and communities. These analyses pinpoint conserved genes that code for traits mediating core cellular functions like transcription and translation, and more variable genes that code for accessory traits like those conferring antibiotic and viral resistance, and metabolic capabilities to name a few [1, 2, 3, 4]. The effects of trait variation established by different and co-occurring genomic regions, on the co-evolution of microbes and their natural enemies, remains a largely unexplored avenue. Of particular relevance to strain diversity are the co-occurring trait axes that respectively confer specificity in ecological interactions with natural enemies, and thus induce density and/or frequency-dependent selection, and variation in intrinsic rates and therefore demography, leading to positive selection. In a purely ecological context, modern coexistence theory establishes the conditions required by the two co-occurring trait axes to give rise to coexistence or exclusion [5]. More recently, work by Good et al. examines similar conditions in a mathematical model for a microbial system with comparable timescales of ecology and evolution where mutations introduce novel trait alleles [6]. The expectations of these previous frameworks do not necessarily apply to high-dimensional trait-spaces with large variation, or to nontrivial network structures of ecological interactions (who interacts with whom) [7, 8]. Moreover, despite advances in addressing coexistence at equilibrium in high-dimensional ecological systems [9, 10, 11], the joint effect of the above co-occurring trait axes also remains poorly understood, especially in systems with comparable timescales of demography and trait innovation. Here, we address this question with a stochastic model for the co-evolutionary host-pathogen dynamics of a microbe-lytic virus system with CRISPR-Cas immune memory.

The CRISPR-Cas system is an immune system found in the accessory genome of many microbial species. This adaptive immune system operates by integrating DNA fragments of infecting viruses, known as ‘protospacers’, into the microbial host’s genome as ‘spacers’ [12]. The CRISPR spacer arrays encoded in the host’s genome thus act as a multi-locus, sequence-specific immune memory of past infections [12]. The presence of a spacer-protospacer match in a subsequent host-virus encounter confers protection against viral infection and lysis. High host and viral strain diversity and non-trivial network interaction structures (‘who infects who’, ‘who is protected from whom’) emerge in the transient temporal dynamics, partly enabled by a large combinatoric trait space from viral repertoire and host memory array sizes, and viral protospacer mutations and host spacer acquisitions. This is in contrast to the simple, one-to-one, infection network structures observed in the ‘kill-the-winner’ model proposed for anti-viral defense mechanisms such as surface resistance and restriction modification systems [13, 14].

Previous studies of CRISPR-mediated co-evolution have largely focused on the emergent and cumulative host immune diversity and structure promoted by negative frequency-dependent selection. In particular, theoretical studies have revealed transient coexistence of host and pathogen, with an alternation of dynamics, between periods when hosts establish control of viral proliferation and those of major viral epidemics with associated rapid host-virus co-diversification. In these transient dynamics, the ultimate fate of the pathogen is extinction. The role of (proto-)spacer diversity and network structure in transitions between these phases have also been extensively addressed [15, 16, 17, 18, 19, 20, 21].

Alongside CRISPR-induced immune memory, competitive abilities for resources can also vary among host strains. Recent short-term co-evolutionary experiments by Guillemet et al. consider a population of *Streptococcus thermophilus* with both CRISPR immune diversity and competitive asymmetries, supporting ‘royal family’ dynamics of host immune strains previously introduced by Breitbart et al. [22, 23]. Namely, after a large viral epidemic the majority of descendent immune strains that fix belong to lineages that are competitively dominant. These competitive asymmetries can induce directional selection among host strains and thus losses of immune diversity, counteracting the diversity-maintaining force of negative frequency-dependent selection previously mentioned. The effect and role of these two co-occurring and opposing modes of selection on CRISPR-mediated co-evolutionary dynamics remain unexplored in theoretical studies to date.

Some previous models have addressed the combined effects of host resistance and competitive differences, for example the classic ‘kill-the-winner’ (KTW) model and its more recent derivatives [13, 14, 24, 25], which emphasize viral predation as the mechanism maintaining the co-existence of hosts with competitive differences. In more general studies of host-pathogen systems, multilocus gene-for-gene models have been used to examine the dynamics emerging from fitness costs associated to the possession of multiple resistance alleles [26, 27]. Current theory has yet to consider the co-evolutionary consequences of host competitive differences in light of the heritable adaptive immunity characteristic of the CRISPR-Cas system.

In this study, we examine the co-occurrence of two key host trait axes: CRISPR-induced memory and competitive asymmetries (see Figure 1 for a schematic diagram). The former yields negative frequency-dependent selection, promoting immune diversity among hosts, while the latter leads to directional selection, which constrains such diversity. We investigate the impact of these counteracting selective pressures on viral evolution and persistence. To this end, we extend a previous computational branching process model of CRISPR-mediated microbe-lytic virus co-evolution to include differences in host competitive abilities [21]. We specifically assume that competitive asymmetries between host strains are manifest as differences in intrinsic growth rates encoded by an underlying trait locus. The intensity of selection acting on this trait determines the breadth of intrinsic growth-rate variation. Total fitness of a host strain is then a function of both its immune memory and intrinsic growth rate. With numerical simulations, we investigate dynamical outcomes of diversity for both host immunity and intrinsic growth rates, systematically examine the effect of increasing host selection intensity on viral persistence and associated metrics, and compare viral adaptation measures for extreme selection regimes. We also discuss correspondences to the empirical observations of Guillemet et al. [22] on ‘royal family’ dynamics.

**Figure 1.**
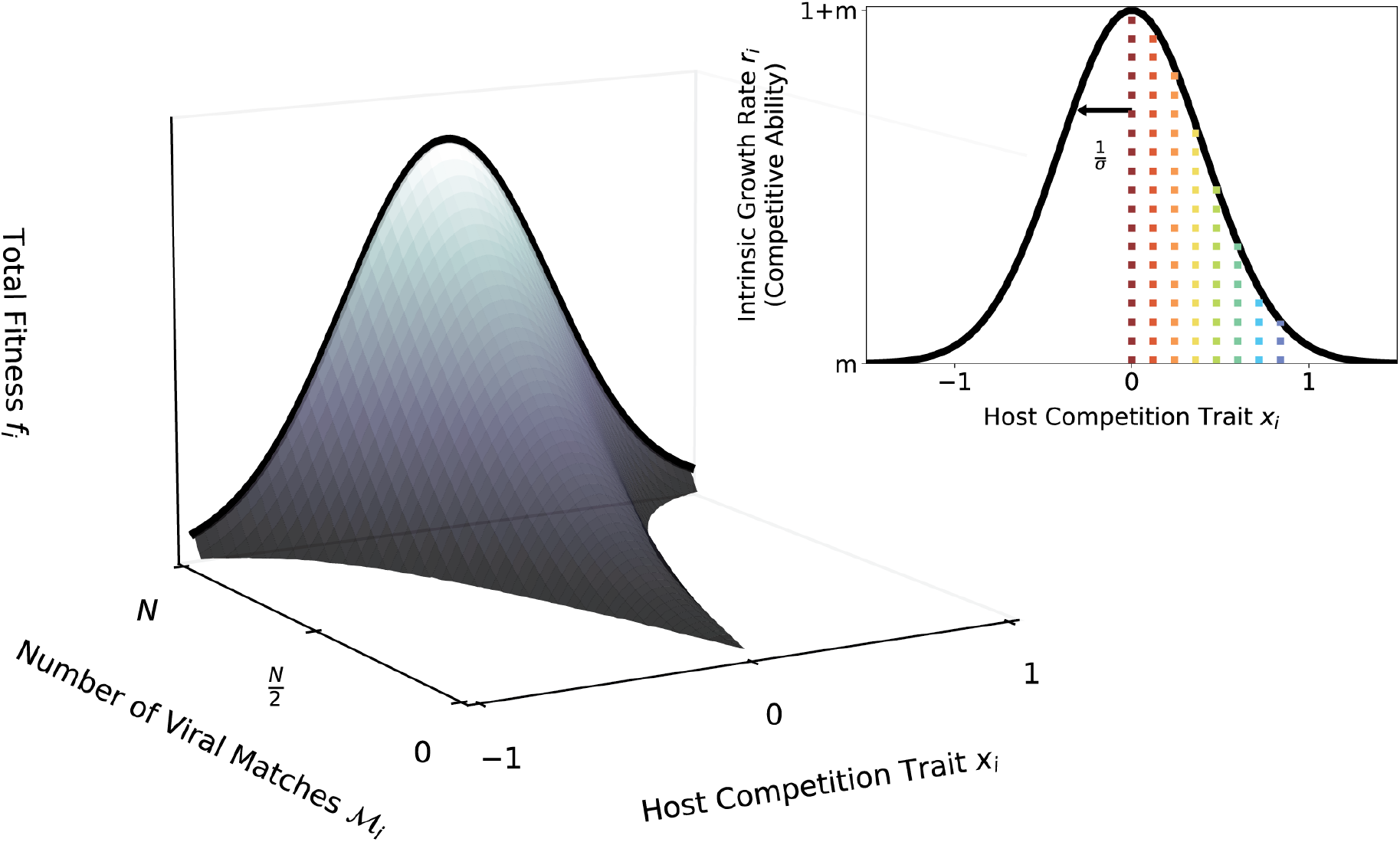
Diagram of host fitness as a function of two trait axes of a microbial host strain. Negative frequency-dependent and directional selection emerge from the variation of such traits, respectively. The axis labeled as host-competition trait has an absolute effect on fitness further depicted in the inset. Namely, the intrinsic growth rates in our model are a continuous Gaussian function of *x*_*i*_, given by 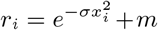, where *m* is the washout rate and *σ* is the associated selection intensity. The associated variation in intrinsic growth rates represents underlying host competitive asymmetries. The other axis represents a component of fitness that depends on the viral diversity and associated frequencies, and is therefore time variable. It monotonically increases with the number of viral matches of the host at a given time. When a host strain matches all viral strains (i.e. ℳ_*i*_ = *N*), or in the absence of viruses, the fitness of a host is dominated by competitive asymmetries leading to directional selection (i.e. competitive exclusion). As ℳ_*i*_ decreases, the host strain becomes susceptible to a larger frequency of the viral population, thus reducing total fitness. The functional form of such decline in the ℳ_*i*_ axis depends on the structure and frequencies of viral diversity, which changes in time. Note that in an extension of the model, the host competition trait randomly mutates upon a spacer acquisition event (with a probability *μ*_*s*_). The colored vertical lines in the inset correspond to the intrinsic growth rates of the example dynamics in Figure (2).

## Methods

### Model

We extend a stochastic model of CRISPR-mediated microbe and lytic virus coevolution, a multitype branching process implemented computationally with a Gillespie algorithm. Events are implemented as an inhomogeneous Poisson process, where time is continuous and event times are exponentially distributed with corresponding rates. Microbial hosts in our model replicate at a rate *r*, and ‘washout’ at a rate *m*. To model asymmetries in competitive abilities, we introduce variation among the intrinsic growth rates *r* of the initialized microbial hosts. We attribute a trait locus to every host strain which encodes an intrinsic growth rate. This locus takes an allelic value *x* from the interval [−1, 1] which encodes for an intrinsic growth rate *r* defined by the continuous Gaussian function *r* = exp(−*σx*^2^) + *m*, where *σ* represents the intensity of selection. As selection intensifies, the half-maximum width of the fitness function shortens, causing the higher fitness values to be represented by fewer trait values around the origin. As selection weakens due to the lack of competitive asymmetries, *σ* → 0, the fitness function approaches uniformity (i.e. neutrality). Figure (1) depicts an example of the map between competition trait alleles and intrinsic growth rates for the 8 host strains used to initialize simulations of our model.

A viral strain naturally decays at a rate *d*, and is defined by a repertoire with a fixed number of loci *g* that carry discrete traits often referred to as *protospacers* [12]. Upon adsorption that occurs at a rate *φ* per particle, a microbe utilizes its CRISPR-Cas immune system to evade lysis with a probability *q*, such that one of the *g* protospacers of a viral strain is randomly selected then integrated as a so-called *spacer* into its genome. The distinct collection of spacers accrued in a microbial host’s lifetime defines their immune type. The spacers confer protection from, and cause the decay of, future viruses that carry at least one matching protospacer. The total fitness of a host strain is thus a function of both its intrinsic growth rate *r* and its spacer array (see Figure 1 for a schematic depiction). Furthermore, recent evidence suggests SNP mutations and INDELs (insertions/deletions) occurring upon spacer acquisition [22, 28]. To investigate possible consequences of such genomic changes, we allow for host traits encoding competitive ability to mutate upon spacer acquisition with a probability *μ*_*s*_. We refer to this as *competitive-ability mutations* hereafter. Moreover, with a probability of 1 − *q* upon adsorption, a virus can successfully lyse a microbe and release a burst of *β* virion daughters, where the probability of having a mutated protospacer is *μ*. Note that we do not assume any variation in the intrinsic demographic rates of the virus, nor any costs from escape mutations (pleiotropic effects). We also note that an infinite allele assumption for viral protospacer mutation is imposed: every protospacer mutation introduces true allelic novelty to the viral population. Here, ‘infinite’ refers to the possible protospacer alleles, as opposed to possible protospacer loci. For corresponding stochastic reactions see Supplementary Information.

We initialize our simulations with a single viral strain and 8 distinct host strains, both represented by 100 individuals. For our primary treatment, *treatment I*, we designate the host strain with the highest intrinsic growth rate to be completely susceptible to the single viral strain, whereas the other seven strains are chosen so that each has a distinct single-spacer match sampled at random from the *g* protospacer loci of a viral repertoire. Alongside, we also consider a second treatment, *treatment II*, comprising of 8 host strains and 8 viral strains. Each viral strain population can infect only one host strain population, where each host strain is protected from other viral strains with a single spacer match. For treatment II, we define the total viral population size to be ~ 100, where each viral strain population is a fraction thereof. Results pertaining to treatment II, and competitive-ability mutations, are in the Supplementary Information.

## Results

### Lineages of host strains with highest competitive abilities gain and maintain dominance, representing the majority of immune diversity

In this section, with numerical simulations of our stochastic model, we establish the long-term co-evolutionary outcomes of diversity in both host adaptive immunity and intrinsic growth rates, emerging from the combined effect of negative frequency-dependent and directional selection. The Gaussian function of Figure (1) exemplifies the distribution of our initialized host strains for selection intensity *σ* = 3 in treatment I. Mueller plots in Figure (2A; see S1 for example of treatment II) depict a realized example of host and virus population dynamics, including forward phylogenies (Figure 2B). Note that, similar to the previous computational model of Liaghat et al. [21], the simulated dynamics exhibit transitions between a regime of sustained host control (SHC) and one of major viral epidemics (MVE). In the SHC regime, the host population is near carrying capacity and small intermittent outbreaks occur. These small outbreaks progressively disassemble the immune structure of the host population, thus giving way to a transition to the MVE regime, where rapid host-virus co-diversification occurs as evident in the phylogenies (see [21] for detailed analysis). Transitions between the SHC and MVE regimes are more clearly observed in Figure (2C), where dynamics are illustrated with linear abundances.

**Figure 2.**
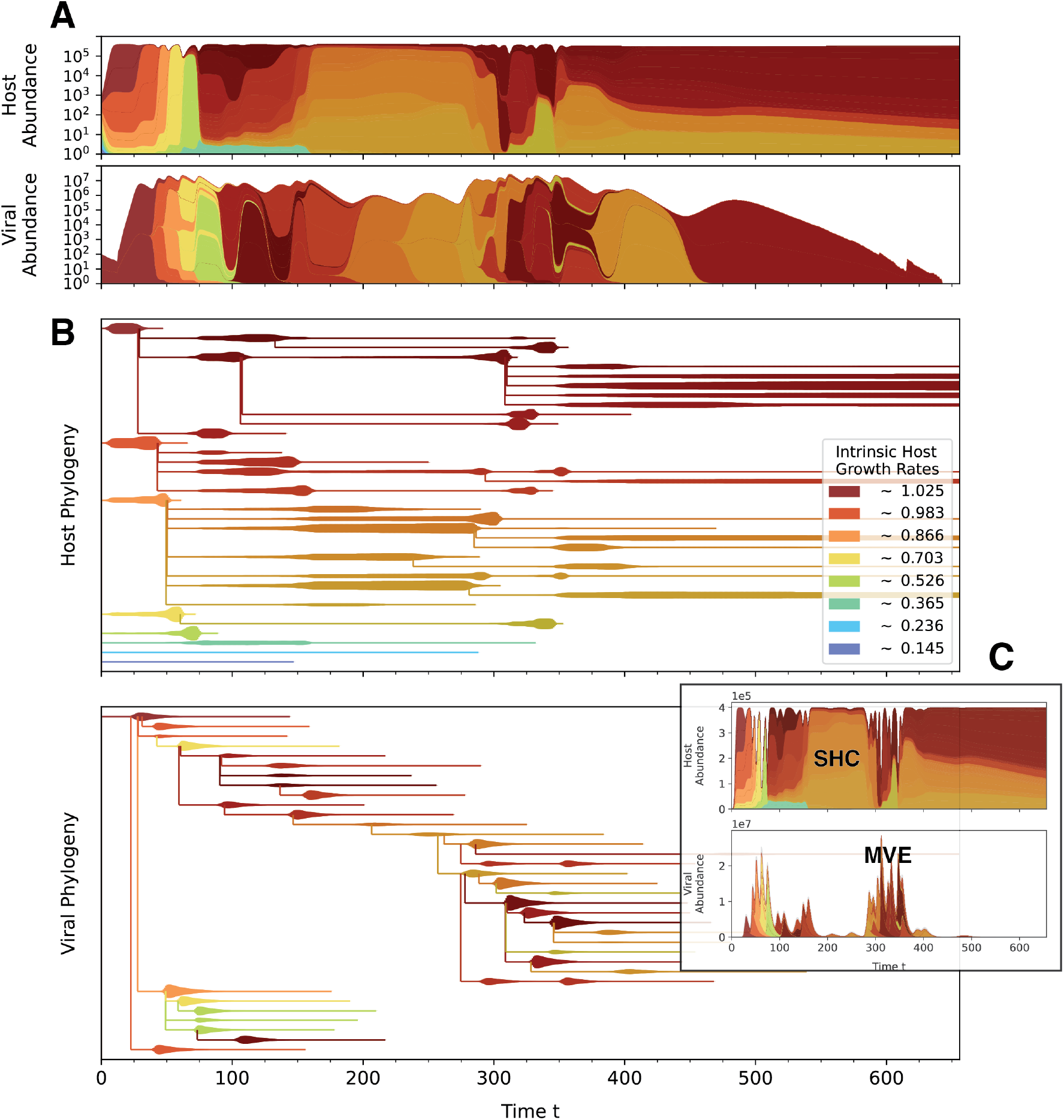
An example simulation of our CRISPR-induced microbe-lytic virus coevolutionary model at a host selection intensity of *σ* = 3 in treatment I. In spite of initial declines due to a major viral epidemic, the most competitively-advantaged host clades regain dominance and diversify the most. In this example, competitive-ability mutations do not occur. See Figure (S1) for example simulation of treatment II. **(A)** Muller plots of host and virus abundances where each stacked color represents the abundance of a respective strain. Total abundances are scaled logarithmically, and distinct strain abundances are scaled linearly. Each hue represents a distinct host clade established by the initial competitors. Initial competitors are represented by a lighter shade of the hue, and its daughter strains are represented by the darker shade. The viral strains are colored with the hue of the most abundant host strain that they infect throughout the entire simulation. **(B)** Forward phylogenies corresponding to (A) where the width of branches represent linearly-scaled abundances of a respective strain. **(C)** Dynamics corresponding to (A) & (B) but represented with linear total abundances. Despite the inclusion of host competitive asymmetries, this model recapitulates the previously described *alternating* dynamics [21].The regime of sustained host control (SHC) is a transient period where the host biomass is saturated at, or near, carrying capacity. The major viral epidemics regime is the short-lived rapid succession of epidemics generated by multiple viral strains, where rapid co-diversification also takes place.

To examine the dominance of the host clades established by the initial competitors, we track the expected mean of the host competitive abilities over time 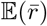. Here 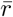 is the mean intrinsic growth rate in a single replicate, and 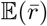 is the expectation of the mean 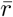 among the 400 simulated replicates. As expected in the absence of viruses, we find that host strains with higher competitive advantages consistently out-compete other strains, causing the mean competitive ability of the host population to gradually converge to the maximal intrinsic growth rate of 1.025 (Figure 3A). However, when viruses are introduced, a rapid decline in the mean competitive ability of the host strains is expected to occur due to a large viral epidemic. Following the rapid initial decline, the mean competitive ability of the host population rebounds to a large value comparable to that of the host population when viruses are absent. For treatment II, Figure (S2B) shows similar rebounding dynamics when a diverse set of viral strains are introduced. These observations are in contrast to KTW models where the dominance of competitors with different intrinsic growth rates cyclically alternate (i.e. fluctuating selection) [24, 13, 14]. Note that the observed resurgence of the dominant competitors remains robust in the case of competitive-ability mutations (Figure S2A; see also treatment II in Figure S2C).

**Figure 3.**
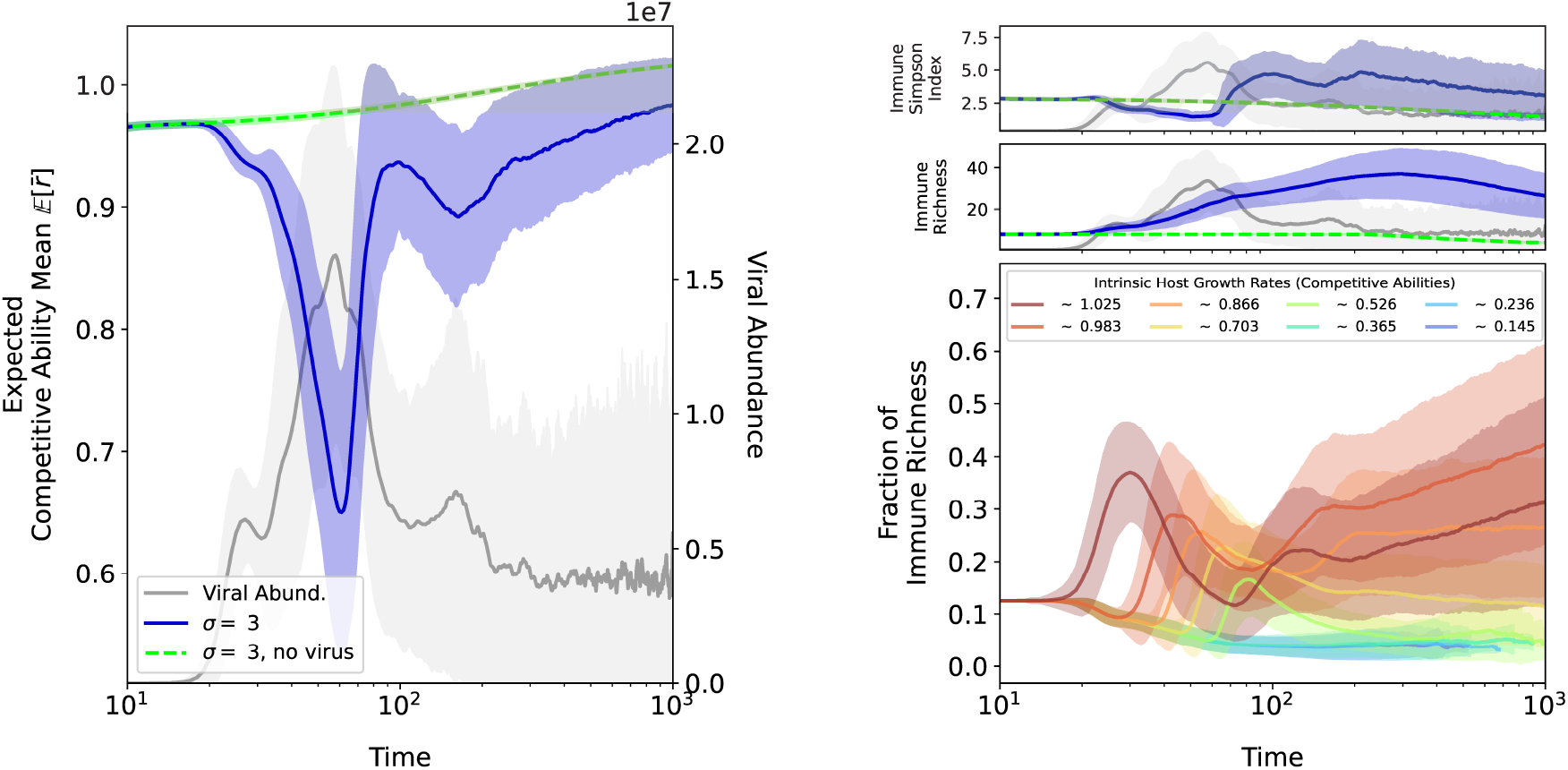
Host diversity expectations computed for a given regime of host selection (*σ* = 3), with and without the viral population (blue and green curves, respectively) in treatment I. Expected total viral abundance is represented by the grey curves. Light shades represent the standard deviation among the 400 simulated replicates. **(A)** The expected competitive ability for the host population over time. ‘Competitive ability’ specifically refers here to the intrinsic intrinsic growth rates of the host strains. 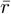 is the mean intrinsic growth rate in a single replicate, and 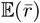 is the expectation of the mean 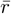 among the 400 simulated replicates. The lighter blue shade represents the standard deviation among replicates, and not within a single replicate. Despite the rapid initial rapid decline of the fittest host strains due to a viral epidemic, the fittest host strains rebound back into dominance. **(B)** The expected Simpson index of the host immune strains over time. Upon the first expected viral epidemic, the Simpson diversity rapidly drops. Thereafter, the Simpson diversity surpasses values of the initial period. **(C)** The dynamics of immune strain richness show that the increase in Simpson diversity is not due to the recovery of older diversity, but rather to the non-stationary diversification that occurs upon immune acquisition. **(E)** The proportion of total immune strain richness represented by the clades established by the initial competitors. The 3 clades with highest intrinsic growth rates diversify the most on average, and thus represent the majority of the immune diversity in the population after the first major viral epidemic.

When the virus is absent, the Simpson index of immune diversity in both treatments gradually declines as expected, due to the competitive exclusion of host strains (Figure 3). The Simpson index is given here by 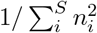 where *n*_*i*_ is the frequency of a host immune strain with a unique spacer array *i*, and *S* is the total number of host immune strains. When the virus is present, this index declines upon the first expected viral epidemic for both treatments I and II. The host phylogenies of Figures (2) & (S1) suggest that this decline of Simpson diversity is accompanied by viral outbreaks that tend to preferentially consume host strains with higher intrinsic growth rates before those with lower intrinsic growth rates. Due to the density dependence of infections, competitively-advantaged host strains which tend to represent larger proportions of the carrying capacity are more likely to be infected than co-occurring competing strains. Upon the termination of the major viral epidemics, the Simpson index rebounds and surpasses levels of the initial period. This rebound of immune diversity is not due to a recovery of the older diversity, but rather to the non-stationary diversification of the host clades. This is demonstrated by the increase in immune richness after the first few viral epidemics in Figure (3B). Here, richness refers to the absolute number of unique spacer arrays in the host population.

Figure (3D) shows the fraction of the immune richness represented by the clades established by the initial competitors. The clades originating from competitively dominant hosts, i.e. those with the highest intrinsic growth rates, diversify the most on average. For our instantiation of host intrinsic growth rates, after the first few viral epidemics, the majority of the immune diversity is represented by 3 clades with the highest intrinsic growth rates (1.025, ~ .983 & ~ .866 in Figure (3D), despite the initial collapse due to viral infections. Consequently, the majority of immune diversity generated from past co-evolution is largely represented by clades established by the initially competitively-dominant host strains. The phylogeny and Muller plot of the host strains in Figure (2) demonstrate an example of how the majority of immune diversification is associated the 3 most competitive clades. In the case of competitive-ability mutations, the immune diversity is represented by a more even distribution of the different host clades. This indicates that the resurgent competitors do not necessarily belong to the clades established by the initially dominant competitors.

### Expected time to ultimate viral extinction increases with asymmetries of host competition

To systematically examine the effect of host selection on viral persistence, we implement a parameter sweep for our model across a range of host selection intensities for both treatments I and II. For each of the treatments, we also consider varied probabilities of competitive-ability mutations. For a geometrically incremented set of host selection intensities (*σ* ∈ [0, 10]), we simulate 400 realizations of the microbe-virus co-evolutionary dynamics. Namely, at one extreme is the regime of no host selection (*σ* = 0), where host strains are competitively equivalent (neutral) and thus have equal intrinsic growth rates, whereas at the other extreme is the regime of strong host selection (*σ* = 10) where competition is strongly asymmetric and introduces variation in the intrinsic growth rates. For each realization, we compute the time to ultimate viral extinction, as well as associated quantities including the number of MVEs, the duration of SHC periods that separate MVEs, the mean number of viral mutants generated and spacers acquired per outbreak.

For our selected parameter values, the viral population eventually goes to extinction. This is a common outcome of CRISPR-mediated co-evolution in previous computational models [20, 21], also observed in experimental studies [29, 30, 31]. Figure (4A) shows that the ultimate viral extinction times exhibit an increasing trend as host selection intensifies, for treatment I in the absence of competitive-ability mutations. A Kruskal-Wallis *H*-test for the trend supports a significant increasing shift in the medians; of extinction times at the higher selection intensities (*p*-value of ~ 6.03 · 10^−68^, see *Supplementary Information* for more details). In correspondence to the increasing trend of extinction times, viral evolution tends to accelerate as a function of host selection intensity. Namely, the viral population more frequently and rapidly overcomes SHC periods, thus generating more MVEs (Figure 4B,C). Note that we designate a major viral epidemic as one that causes a large decline of the host population (of *at least* 45%). Furthermore, as host selection intensifies, the mean number of viral mutants generated per outbreak increases, whereas the mean number of spacers acquired per outbreak decreases (Figure 4C & D). This suggests that a decline in host immune diversity, driven by strong directional selection, facilitates a disproportionate increase in viral density and thus diversity. Such viral diversification increases the likelihood of escapes from host immunity. In the case of treatment II, similar to treatment I, an increasing trend of time to ultimate viral extinction is observed as host selection intensifies. Increasing and decreasing trends for the number of MVEs and duration of SHC periods, respectively, are also observed. However, in contrast to treatment I, the trends appear to converge as host selection intensifies and exhibit larger variance (Figure S4A). This may be an effect induced by the drift of viral strain populations. Each viral strain population in treatment II is represented by a small fraction of 100 individuals (Methods), and consequently is subject to more stochasticity than a single viral strain population in treatment I. The emergence of an epidemic by a given viral strain is therefore less likely in treatment II, than in treatment I. An extensive exploration of the effect of inocula sizes, and its associated stochasticity, is outside the scope of this work and remains for future studies.

**Figure 4.**
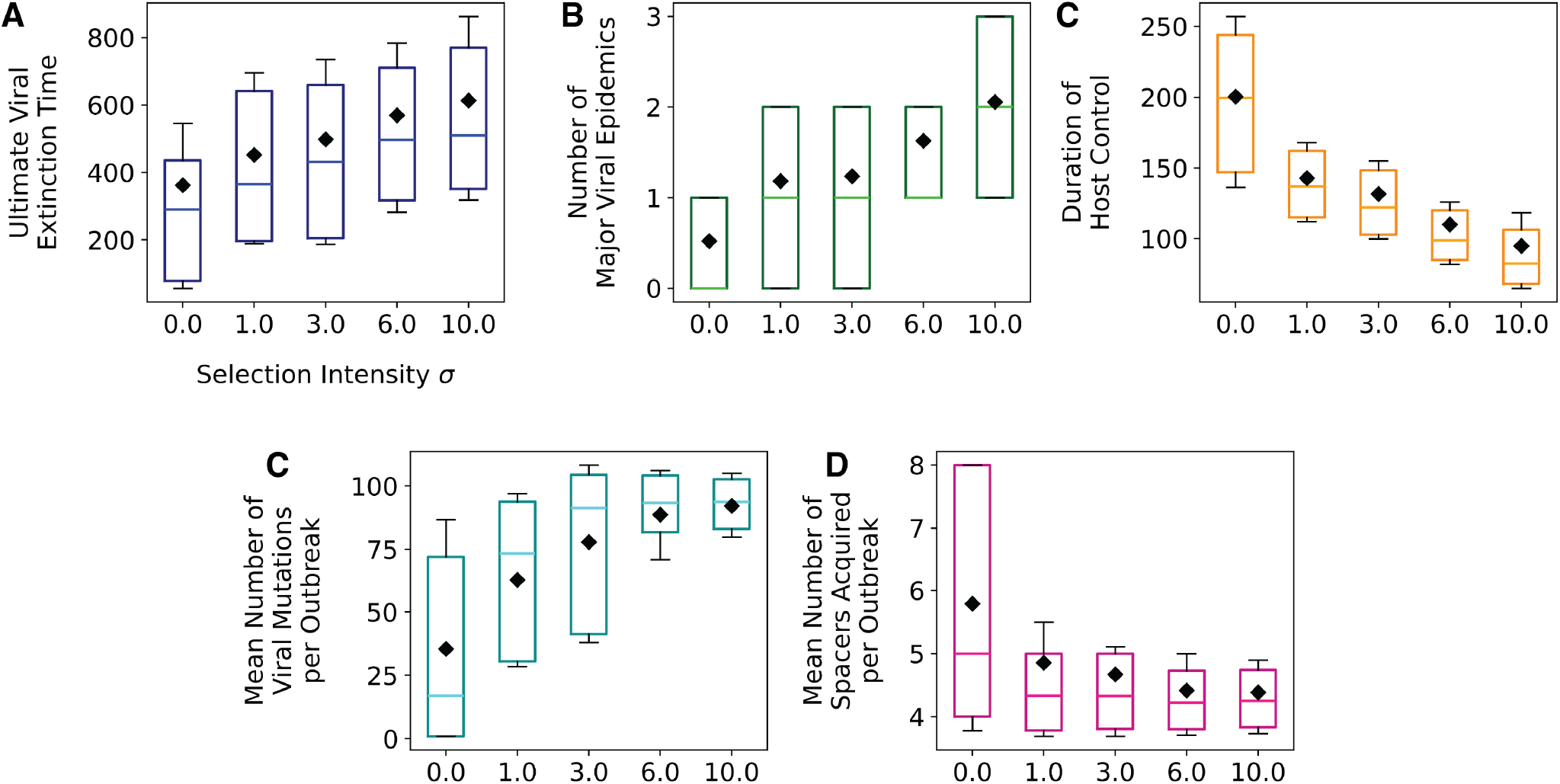
Box-and-whisker plots capture distributions of the time to ultimate viral extinction, and associated metrics, as a function of host selection intensity *σ* in treatment I. The inter-quantile ranges represent 20-80% of the simulated replicates. The black diamonds represent mean values, and the colored horizontal lines in the inter-quantile ranges represent medians. **(A)** As host selection intensifies, times to ultimate viral extinction tend to increase (*p* ~ 6.03·10^−68^ from a Kruskal-Wallis *H*-test), **B** major viral epidemics also become more frequent. **(C)** The duration of host control, which transiently separate these major viral epidemics, also shortens. This pattern suggests rapid disassembly of host immune structure. **C** The mean number of viral mutants per outbreak also tends to increase as host selection intensifies, whereas the mean number of spacers acquired per outbreak decreases **D**. This suggests that recurrent viral escapes are more likely as host selection intensifies. See Figure (S3) for emergent trends in the context of competitive-ability mutations, and Figure (S4) for treatment II.

Moreover, the increasing trend in viral extinction time is lost as competitive-ability mutations become more probable in both treatments I and II (Figure S3.1,S4.1). Despite this weakening of the trend, viral evolution tends to accelerate as host selection intensifies. This is demonstrated by the increase in MVEs and shortening of SHC durations as host selection intensifies, which also become concentrated into the earlier times of the dynamics (Figures S3.3-5, S4.3-5). Furthermore, as competitive-ability mutations become more probable, the mean number of spacers acquired per outbreak reveals an increasing trend as a function of host selection intensity (Figure S3.7 & S4.7). This is in contrast to the decreasing trend observed in the absence of competitive-ability mutations (Figure 4D), suggesting that with the acceleration of viral evolution, host immune evolution also accelerates, promoting the rapid assembly of immune structure needed to extinguish the viral population.

During periods of sustained host control, a viral strain consumes virtually all of its pool of susceptible host strains upon a small outbreak [21]. Therefore in order for the total viral population to adapt and further persist, a viral strain must escape competing host strains upon an outbreak. To obtain a closer look at the effect of host selection on viral escape over time, we next consider two summary quantities often used to investigate co-evolutionary dynamics, namely temporal and local adaptation.

### Strong competitive asymmetries of host population promotes viral evolution and adaptation

Each viral escape permits access to a new pool of susceptible host strains, which increases the fitness of the total viral population. This effect on viral fitness however is only transient, as the viral population tends to *burn* through its pool of susceptible host strains (see [21]). If the viral population can generate another escape variant upon an outbreak, the total viral population can regain fitness and persist for another duration of time. Along with viral escapes, the host can acquire spacers upon an outbreak. This acquired immunity also tends to accumulate during host control periods. The long-term persistence of the viral population is thus contingent on its ability to recurrently escape acquired immunity and consequently generate outbreaks. Figure 4 suggests that host selection intensity must then modulate these recurrent, and temporally localized, gains and losses of viral fitness.

We examine changes of viral fitness more closely in the context of different host selection intensities, with measures of viral temporal adaptation (TA) and local adaptation (LA). Both quantities are functions of mean viral fitness 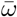 (SI). The temporal adaptation quantity in Equation (S1) captures on average how adapted the viral population is to the host population from a retarded or advanced time shift of *τ*. This quantity can be used in empirical or experimental settings to identify the type of selection occurring in a co-evolutionary system [32, 33]. The local adaptation quantity in Equation (S2) determines whether the virus is, on average, more adapted to sympatric (same replicate) than to allopatric (different replicate) host populations. See Equations (S1) & (S2) in SI for full expressions of TA and LA.

We observe that viral TA to the host population is maximal in the recent past, and exhibits a decreasing trend, for both extrema of host selection intensity: *σ* = 0 & *σ* = 10 (see Figures 5 and S5 for treatments I and II, respectively). This decreasing trend reflects that the viral population overcomes accumulated host immunity through recurrent escapes throughout the dynamics. In contrast, viral TA rapidly drops when matched against hosts from the future. This demonstrates how rapidly hosts are able to acquire spacers, and thus immunity, from dominant viral strain. Furthermore, in the context of strong host selection, viral TA in the recent past is steeper in slope than in the absence of host selection. This corresponds to the acceleration of viral evolution suggested by the previous trends of the number of major viral epidemics, the durations of the SHC period in between, and their times of occurrence (Figures 4, S3, S4). The observed form of viral TA also suggests ‘arms-race’ frequency dynamics of viral diversity [32, 33]. The augmented slope in the recent past is also observed in the context of competitive-ability mutations. Despite the detrimental effect of competitive-ability mutations on viral persistence, this result is consistent with the observed acceleration of viral evolution observed in Figures (S3),(S4).

**Figure 5.**
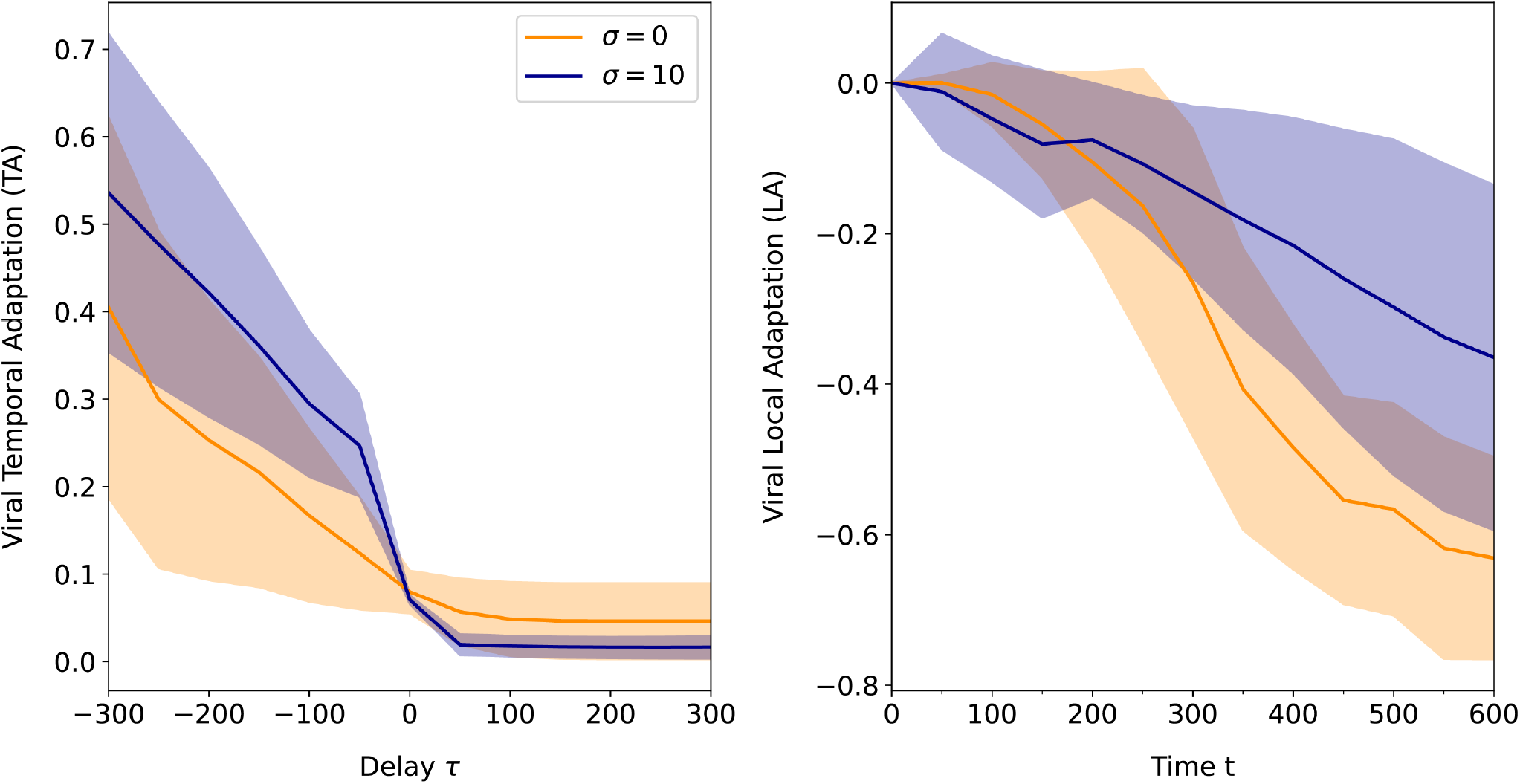
Temporal adaptation (TA) and local adaptation (LA) measures of the viral population for both extrema of host selection intensity (*σ* = 0 & *σ* = 10). The temporal adaptation quantity captures on average how adapted the viral population is to the host population from a retarded or advanced time shift of *τ*. The local adaptation quantity determines whether the virus is on average more adapted to sympatric (same replicate) than to allopatric (different replicate) host populations. Here viral adaptation refers to the mean viral fitness, i.e. mean frequency of susceptible hosts per viral strain. See Equations (**??**) & (**??**) for more details. Note that TA = 1 indicates that the viral population is on average completely adapted to the host population of a time shift *τ*, and TA = 0 indicates the viral population is completely maladapted. Also note that LA = 1 indicates that the viral population is completely sympatrically adapted but not allopatrically adapted, whereas LA = −1 indicates the reverse. TA demonstrates a declining trend for both selection intensities, which indicates the propensity for viral maladaptation. Both TA and LA rapidly drop for both extrema of host selection intensity. However, in the absence of host selection, viral maladaptation is more rapid than for strong selection.

Lastly, we observe that viral local adaptation (LA) rapidly declines for both cases of host selection intensity (Figure 5 and S6 for treatments I and II, respectively). In particular, we observe that LA declines more slowly in a regime of strong host selection intensity, regardless of competitive-ability mutations (Figure S6). This implies that, on average, the viral population evolves to be more sympatrically adapted in the context of strong host selection than in the absence host selection. Interestingly, local mean viral fitness for both cases of selection are expected to remain relatively constant, and of comparable values, throughout the dynamics (Figure S7). The mean viral fitness among contemporaneous, allopatric host populations is thus the main driver of the observed differences of LA in the two cases of host selection. Namely, allopatric adaptation is more rapid in the absence of host selection.

## Discussion

By including host competitive asymmetries in a previous stochastic model of CRISPR-induced microbe-lytic virus co-evolution, we are able to gain insight into the effects of two co-occurring modes of selection on viral evolution. These modes correspond respectively to negative frequencydependent selection arising from immune microbial memory, and directional selection from host competitive asymmetries. Following an initial major decline in host abundance due to a viral epidemic, competitively advantaged host strains rebound to maintain dominance in the population, representing the majority of immune diversity generated throughout the co-evolutionary dynamics. As host selection intensifies, the time to ultimate viral extinction increases which is also associated to the acceleration of viral evolution. Temporal and local adaptation measures also document short and long term behaviors of viral (mal-)adaptation in the two extreme regimes of host selection.

Our numerical results in conjunction with previous experimental work [22], support the recent ‘royal family’ hypothesis [23], which posits that newly rising host genotypes are likely to descend from previous genotypes that are dominant due to intrinsic asymmetries in competitive abilities. In ‘royal family’ dynamics, host strains with competitive advantages maintain dominance despite viral predation. This is in contrast to ‘kill-the-winner’ dynamics, where the preferential targeting of host strains with competitive differences, by the viral populations, alternates between those with high and low competitive advantages. Each host strain thus undergoes cycles of high and low frequencies, reflecting fluctuating selection [13, 14, 24, 25]. Specificity in such preferential targeting would apply to surface resistance and restriction modification systems. It remains an open question whether these two different types of frequency-dependent dynamics arise due to memory operating at different organizational levels: both the individual and population level for CRISPR-induced immunity in royal family, and solely the population level for kill-the-winner. Also, despite the frequent co-occurrence of these differing lines of viral defense mechanisms in natural microbial populations, theoretical expectations of resulting eco-evolutionary dynamics are sparse and remain open for future research (for examples of multi-defense in general host-parasite systems see [34, 35]). Furthermore, ‘royal’ viral lineages may also emerge due to competitive differences from variation in other demographic and interaction parameters in both microbes and viruses.

The observed trend of viral temporal adaption in our study is consistent with ‘arms-race’ frequency-dependent dynamics among viral strains [32, 33] and with viral temporal adaptation signatures in both the monomorphic and polymorphic experimental treatments of Guillemet et al. [22]. The characteristic punctuated nature of viral and host diversification in ‘arms race’ dynamics, and its associated periods of explosive diversification, are so far unique to CRISPR-mediated coevolution. Moreover, viral local adaptation demonstrates a declining trend over time for both of our treatments I and II. In the monomorphic treatments of Guillemet et al., a partial decline may be observed as the experiments terminate, but the significance of this decline remains inconclusive. Our model assumes ‘infinite protospacer alleles’ (Methods), where every mutation introduces a novel protospacer allele among the *g* protospacer loci that define a viral strain. It therefore applies to the scenario of rapid loss of deleterious protospacer alleles occurring from back-mutations. The resulting trend of viral local adaptation captures potential long-term co-evolutionary behavior of empirical systems. In contrast, the experiments of Guillemet et al. capture short-term co-evolutionary behavior. Future experiments examining the long-term behavior of CRISPR-mediated co-evolutionary dynamics, will aid in further corroboration of our theoretical expectations. In addition, molecular experiments examining the probability of protospacer back-mutations will help refine how viral diversification is modeled.

Previous studies have demonstrated the roles of both absolute viral and host abundance in the stochastic emergence and escape of rare viral strains. Chabas et al. demonstrate the positive, non-linear relationship between the size and probability of emergence of initial viral inocula [19]. In parallel, Liaghat et al. show that viral strain escape and emergence is limited by the intensity of host density-dependent competition [21]. In particular, strong density-dependent competition (parameterized by the host’s carrying capacity) limits the total possible size of susceptible hosts through which a rare viral strain can generate outbreaks and thus diversify. Therefore, despite possible increases of the probability of emergence through the augmentation of viral inocula sizes, a decrease in the carrying capacity can counteract this effect. Synthetic empirical and theoretical studies exploring the combined effects of initial viral inocula sizes, and host density-dependent and asymmetric competition, on the probability of viral emergence and long-term persistence are still missing.

Our results indicate that host competitive asymmetries can facilitate viral escape, thus delaying total viral extinction. Nevertheless, the dynamics of our computational model still support the strong propensity for total viral extinction and the consequent transience of the co-evolutionary dynamics described in previous computational and experimental studies [21, 20, 18, 29, 30, 31]. In natural microbe-virus communities, lytic viruses persist and the trailer-end spacers of host CRISPR-Cas arrays appear conserved for approximately 5 years or longer [36, 37]. The question of viral persistence thus requires further consideration, with a metapopulation context being one such direction. Notably, the measure of viral local adaptation in this study reveals strong allopatric adaptation, suggesting that viral emigration to foreign localities may indeed prolong viral persistence.

As microbial pangenomics further uncover co-occurring trait types defining ecological interactions as well as their associated timescales of evolution in natural communities, a deeper understanding of how associated selective forces combine to affect co-evolutionary dynamics and intra- and inter-specific diversity, is becoming possible. Future theoretical studies can define expectations for co-evolutionary and structural patterns in natural communities. Our study takes a step in this direction by showing that competitive asymmetries of hosts can alter the timescale of co-evolution that results from purely negative frequency-dependent selection.

## Acknowledgements

This work was supported by the Gordon and Betty Moore Foundation, GBMF 9195, https://doi.org/10.37807/GBM and by the GEMS Biology Integration Institute, funded by the National Science Foundation DBI Biology Integration Institutes Program, Award Number 2022049.

