## Supplementary Information for "Host Competitive Asymmetries Accelerate Viral Evolution in a Microbe-Virus Coevolutionary System"

### Kruskal-Wallis $H$-test

This statistical test is the non-parametric equivalent of a one-way analysis of variance (ANOVA). Namely, the null hypothesis of this test is that each sample originates from the same distribution. This test determines whether at least one sample stochastically dominates one other sample, i.e. an increased shift in at least one of the sampled distributions. This shift is often associated with a shifted median, especially if the shapes of the sampled distributions are equivalent. Note that we use a non-parametric statistical test due to the long tails of the distributions in Figure (3A) in the main text.

### Temporal and Local Adaptation

Here we define mean viral fitness as $\bar{\omega} = \sum_i h_i(t)v_i(t)$ where $v_i(t)$ denotes the frequency of a viral strain $i$ at a time $t$, and $h_i(t)$, the frequency of hosts susceptible to a viral strain $i$ at time $t$. The quantity of temporal adaptation (TA), for a retarded or advanced timeshift of $\tau$ and a replicate $r$ is given by

$$TA(\tau, r) = \frac{1}{n_t - |\tau|} \sum_{t=\max(0,-\tau)}^{\min(n_t, n_t-\tau)} \sum_{i=1}^{n} h_i(r, t+\tau) p_i(r, t) \tag{S1}$$

where $n_t$ is the total number of sampled time points of a replicate. This quantity captures how much, on average, the viral population has, or will have, become (mal-)adapted to the host population after an elapsed time (delay) $\tau$. Furthermore, the quantity of local adaptation (LA) for a time point $t$ and replicate $r$ is given by

$$LA(r, t) = \sum_{i=1}^{n} h_i(r, t) p_i(r, t) - \frac{1}{n_r - 1} \sum_{j \neq r} \sum_{i=1}^{n} h_i(i, j) p_(j, t) \tag{S2}$$

where $n_r$ is the number of replicates. This quantity captures whether the viral population is more or less sympatrically adapted (to the host population in the same replicate) than it is allopatrically adapted (to the host population in other replicates). A value of 0 indicates equivalence of sympatric and allopatric adaptation.

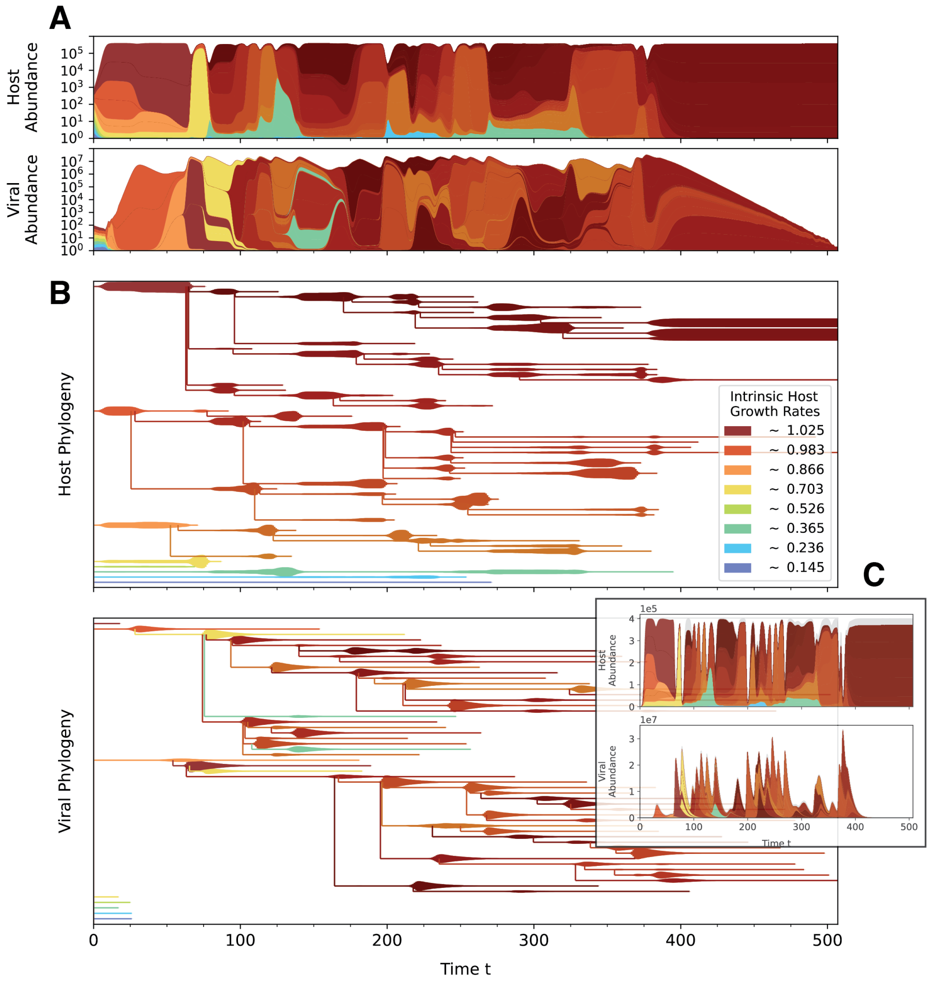


Figure S1: An example simulation of our CRISPR-induced microbe-lytic virus coevolutionary model at a host selection intensity of $\sigma = 3$ in treatment II. In this example, competitive-ability mutations do not occur. **(A)** Muller plots of host and virus abundances where each stacked color represents the abundance of a respective strain. Total abundances are scaled logarithmically, and distinct strain abundances are scaled linearly. Each hue represents a distinct host clade established by the initial competitors. Initial competitors are represented by a lighter shade of the hue, and its daughter strains are represented by the darker shade. The viral strains are colored with the hue of the most abundant host strain that they infect throughout the entire simulation. **(B)** Forward phylogenies corresponding to (A) where the width of branches represent linearly-scaled abundances of a respective strain. **(C)** Dynamics corresponding to (A) & (B) but represented with linear total abundances. Despite the inclusion of host competitive asymmetries, this model recapitulates the *alternating* dynamics addressed in [**?**].The regime of sustained host control (SHC) is a transient period where the host biomass is saturated at, or near, carrying capacity. The major viral epidemics regime is the short-lived rapid succession of epidemics generated by multiple viral strains, where rapid co-diversification also takes place.

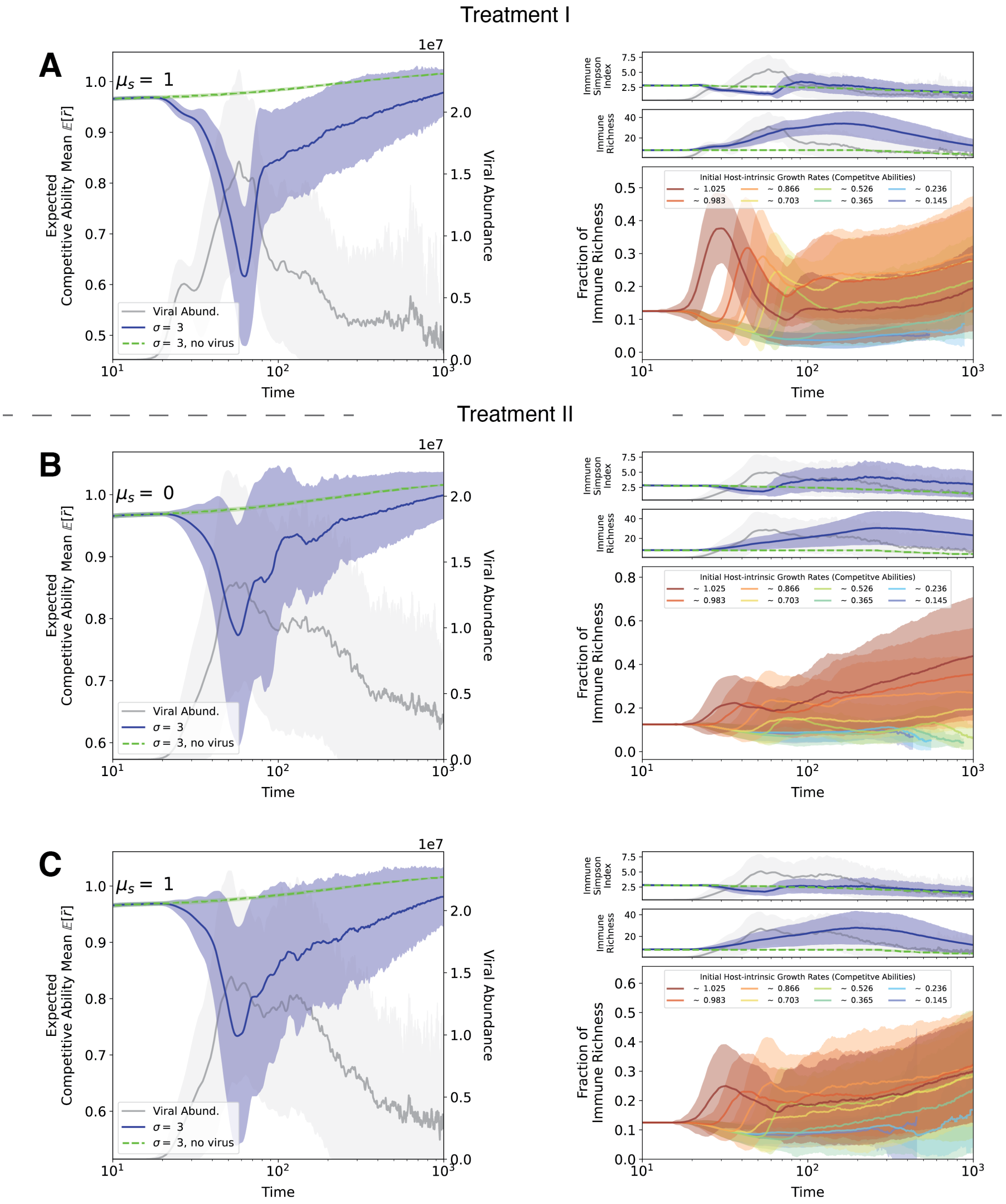


Figure S2: Host diversity expectations computed for a regime of host selection ($\sigma = 3$), for instances with and without the viral population (blue and green curves, respectively) in **(A)** treatment I in the context of competitive-ability mutations $\mu_s = 1$, and treatment II **(B)** with and **(C)** without competitive-ability mutations ($\mu_s = 0$ and $\mu_s = 1$, respectively). Expected total viral abundance is represented by the grey curves. Light shades represent the standard deviations among the 400 simulated replicates. Note that $\bar{r}$ is the mean competitive ability, i.e. intrinsic growth rate, of the host population within a single replicate. The expected competitive ability mean $\mathbb{E}(\bar{r})$ for the host population over time for all treatment show that the fittest host strains rebound back into dominance despite their rapid initial decline due to a viral epidemic. Upon the first expected viral epidemic, the expected Simpson index of the host immune strains over time drops, and thereafter either reaches or surpasses values of the initial era. The expected immune strain richness show that the rebound in Simpson diversity is not due to the recovery of older diversity, but rather to the non-stationary diversification that occurs upon immune acquisition. In the absence of competitive-ability mutations **B**, the clades with the highest intrinsic growth rates diversify the most on average, and thus represent the majority of the immune diversity in the population after the first major viral epidemic. However, when competitive-ability mutations are introduced, the clades established by hosts with lower demographic are able to diversify more and thus represent a larger proportion of the immune diversity after a major viral epidemic.

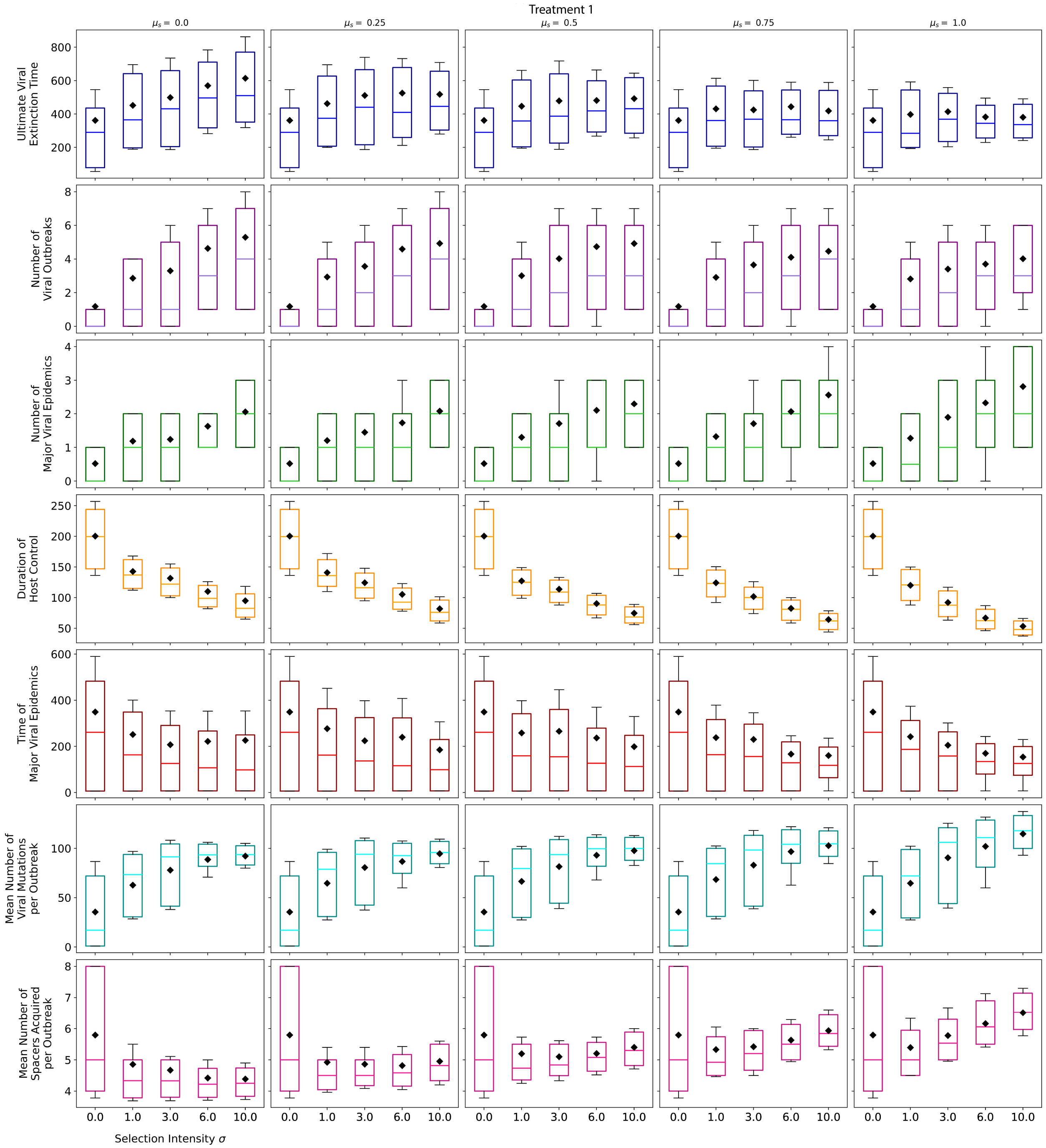


Figure S3: See caption in Figure S4 for summary.

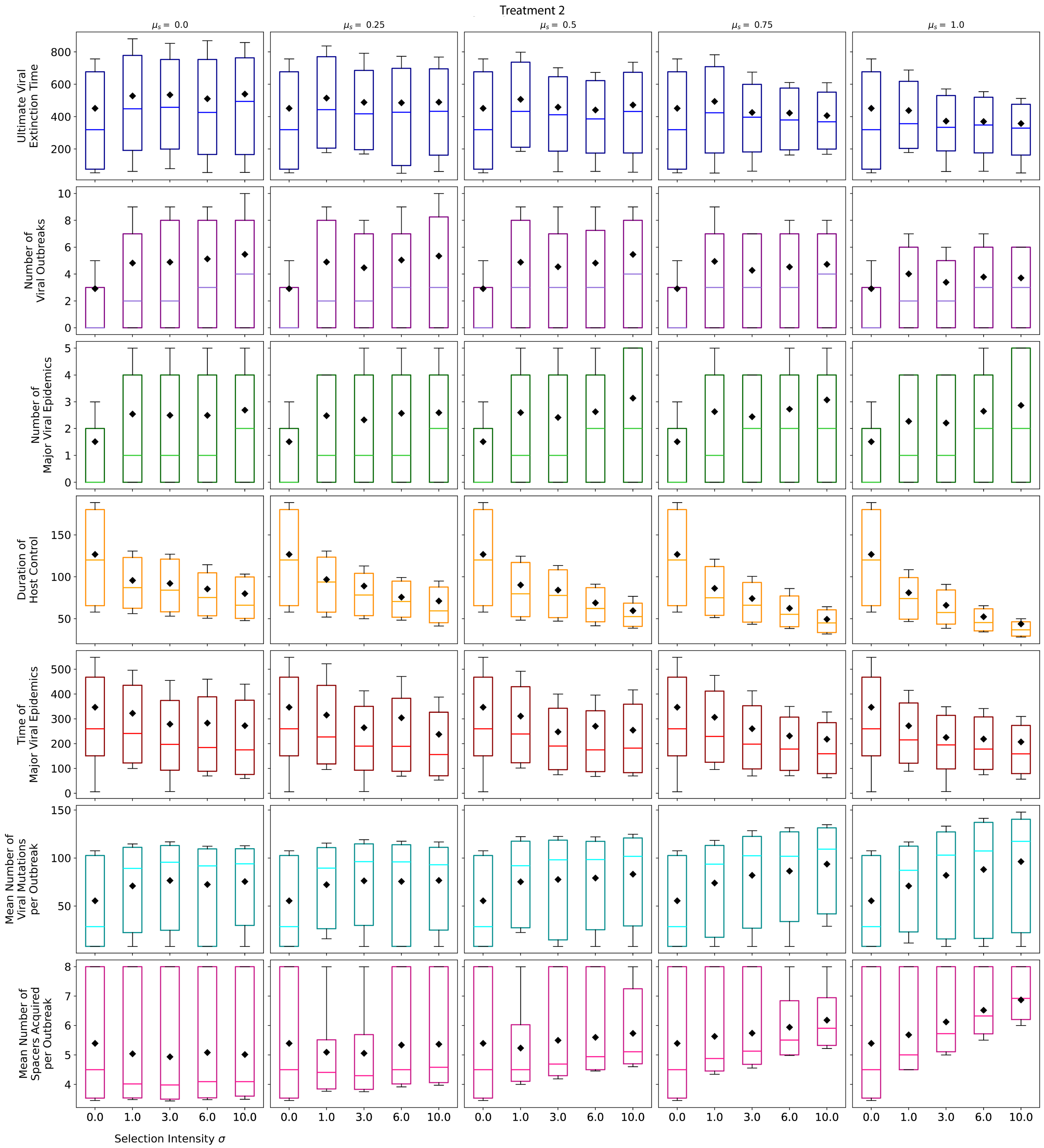


Figure S4: Box-and-whisker plots capture distributions of quantities capturing key features of host-virus dynamics as a function of host selection intensity $\sigma$ in both treatments **(Figure S3)** I and **(Figure S4)** II with varied competitive-ability mutation probabilities. The inter-quantile ranges represent 20-80% of the simulated replicates. The black diamonds represent mean values, and the colored horizontal lines in the inter-quantile ranges represent medians. **(1)** As host selection intensifies in the absence of spacer-induced mutations, times to ultimate viral extinction tend to increase (Kruskal-Wallis $H$-test gives $p \sim 6.03 \cdot 10^{-68}$ for treatment I and $p \sim 1.63 \cdot 10^{-4}$). However, with the increase of the competitive-ability mutation probability, the increasing trend of ultimate viral extinction times is lost. **(2)** The number of outbreaks that consume at least 15% of the host population increase as host selection intensifies. However, the trend weakens as the competitive-ability mutation probability increases. **(3)** The number of major viral epidemics — outbreaks that consume at least 45% of the host population — increases as host selection intensifies. This trend further increases with the probability of competitive-ability mutations. **(4)** The duration of the host control periods that separate the major viral epidemics shorten as host selection intensifies, and further shorten with an increase in competitive-ability mutation probability. **(5)** The alternations between control period and major epidemics occur earlier in the dynamics as host selection intensifies, and occur even sooner as the competitive-ability mutation probability increases. **(6)** The mean number of viral mutants per outbreak tends to increase as host selection intensifies, **(7)** whereas the mean number of spacers acquired per outbreak decreases. However, as the competitive-ability mutation probability increases, the mean number of spacers acquired per outbreak transitions from a declining trend into an increasing one.

### Stochastic Reactions of Model

This system consists of virions $V$ of strain types defined by a set of integers $\rho$, where each integer represents a unique 'protospacer' allele and the repertoire length is fixed, i.e. $|\rho| = g$. These virions infect microbial species $B$ of strain types defined by a set of integers $\alpha$, where $|\alpha| \in [0, \sigma]$ and $\sigma$ represents the host's spacer capacity. Here, $V_\rho$ and $B_\alpha$ represent individuals and not abundances, indexed by the sets that define their respective strain identities. In our model, we impose an infinite allele assumption for viral protospacer mutation — i.e., every mutation introduces true allelic novelty to the viral population.

A microbial host in our model successfully replicates with a probability $r_\alpha(1 - \frac{N}{K})$, where $\frac{r_\alpha N}{K}$ captures the effect of competition,

$$B_\alpha \xrightarrow{r_\alpha(1-N/K)} B_\alpha B_\alpha \tag{S3}$$

$N$ is the total host abundance and $K$ is the carrying capacity. Note that we are assuming similar timescales of host replication and competition in our model. Upon adsorption that occurs at with a probability of $\varphi$, microbes utilize their CRISPR-Cas immune system to evade lysis with a probability $q$, such that one viral protospacer is randomly integrated as a 'spacer' into their respective genomes,

$$B_\alpha V_{\rho \cap \mu = \varnothing} \xrightarrow{\varphi q} B_{\mu'} \tag{S4}$$

where $|\mu' \cap \rho| = 1$. The distinct collection of spacers accrued in a microbial host's lifetime defines their immune type and their memory of previous infection. Namely, the spacers confer protection from, and cause the decay of, future viruses that carry at least one matching protospacer,

$$B_\alpha V_{\rho \cap \mu \neq \varnothing} \xrightarrow{\varphi} B_\alpha \tag{S5}$$

where $|\rho \cap \mu| \geq 1$. Alternatively upon an adsorption, a virus can successfully lyse a microbe and release a burst of $\beta$ virion replicates with a probability of $1 - q$

$$B_\alpha V_{\rho \cap \mu = \varnothing} \xrightarrow{\varphi(1-q)} \underbrace{V_\rho \ldots V_\rho}_{\beta \text{ virions}} \tag{S6}$$

Note that with probability $1 - (1 - \mu)^g$ one of the $\beta$ virions mutates from a strain $\rho$ to $\rho'$, where the probability of protospacer mutation per locus site is $\mu$ and the total repertoire size is $g$. Also, a virus decays with a probability $d$

$$V_\rho \xrightarrow{m} \emptyset \tag{S7}$$

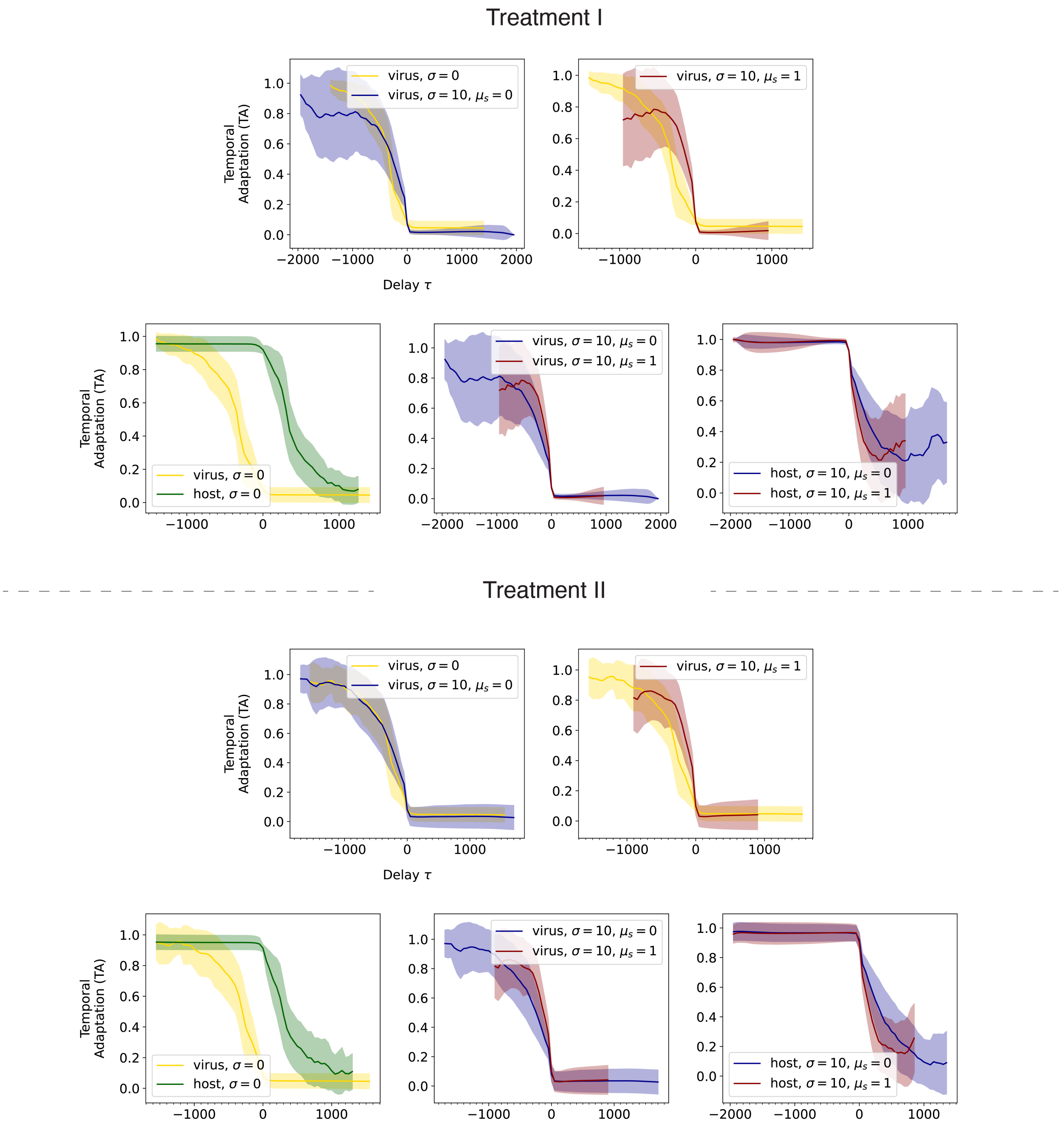


Figure S5: Viral temporal adaptation (TA), along with host mean resistance, for both extrema of host selection intensities ($\sigma = 0$ and $\sigma = 10$) in treatments I and II, and in the presence and absence of competitive-ability mutations. Signature of viral temporal adaptation resembles those derived from experimental results of Guillemet et al. [**?**]. In the case of strong host selection intensity, slope of viral TA near $\tau = 0$ is much steeper than that of viral TA in the absence of host selection. This slope further steepens in the presence of competitive-ability mutations.

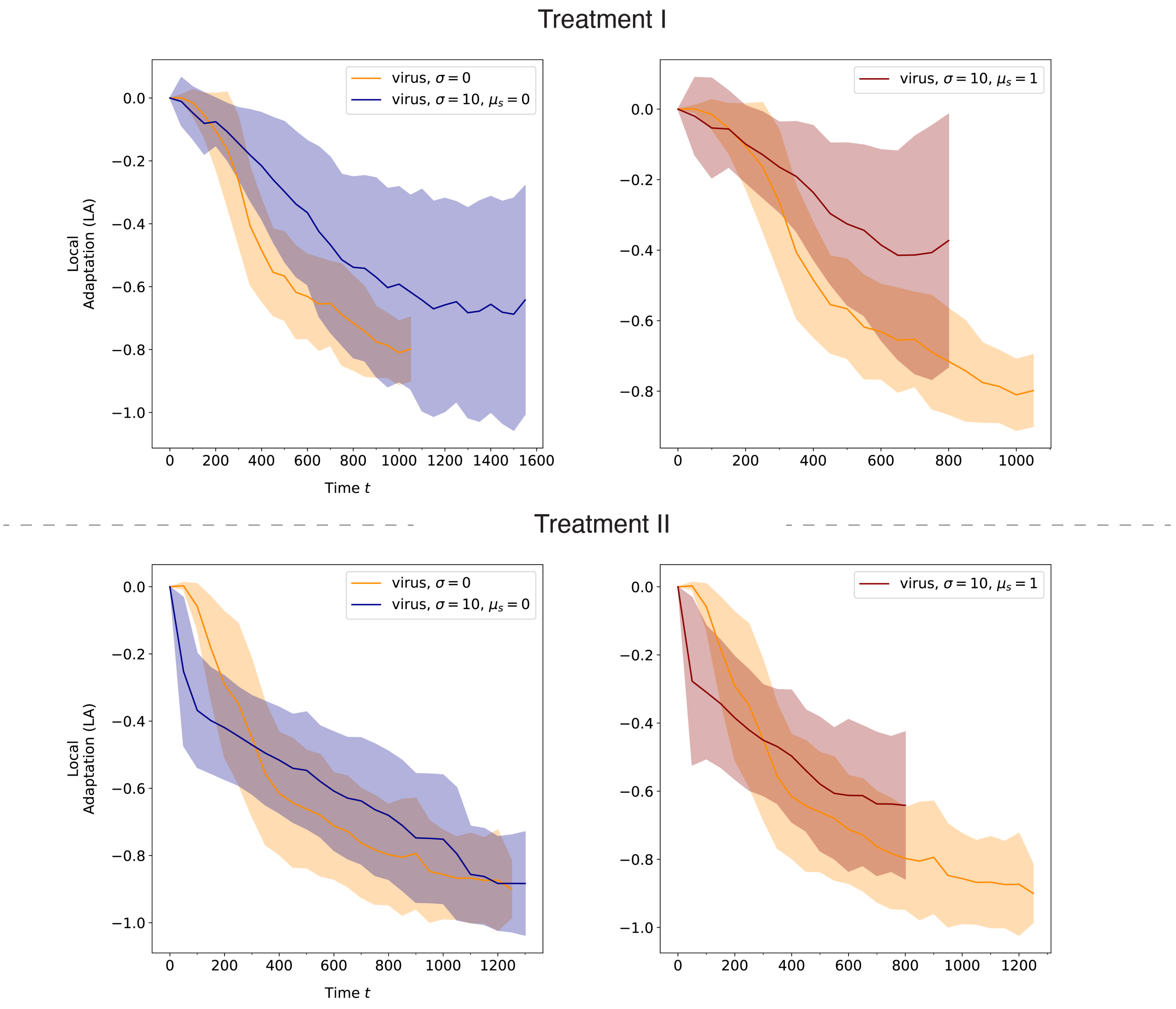


Figure S6: Viral local adaptation (LA) for both extrema of host selection intensities ($\sigma = 0$ and $\sigma = 10$) in treatments I and II, and in the presence and absence of competitive-ability mutations. LA in the absence host selection tends to be lower than that of strong host selection, for all both instances of competitive-ability mutation probabilities.

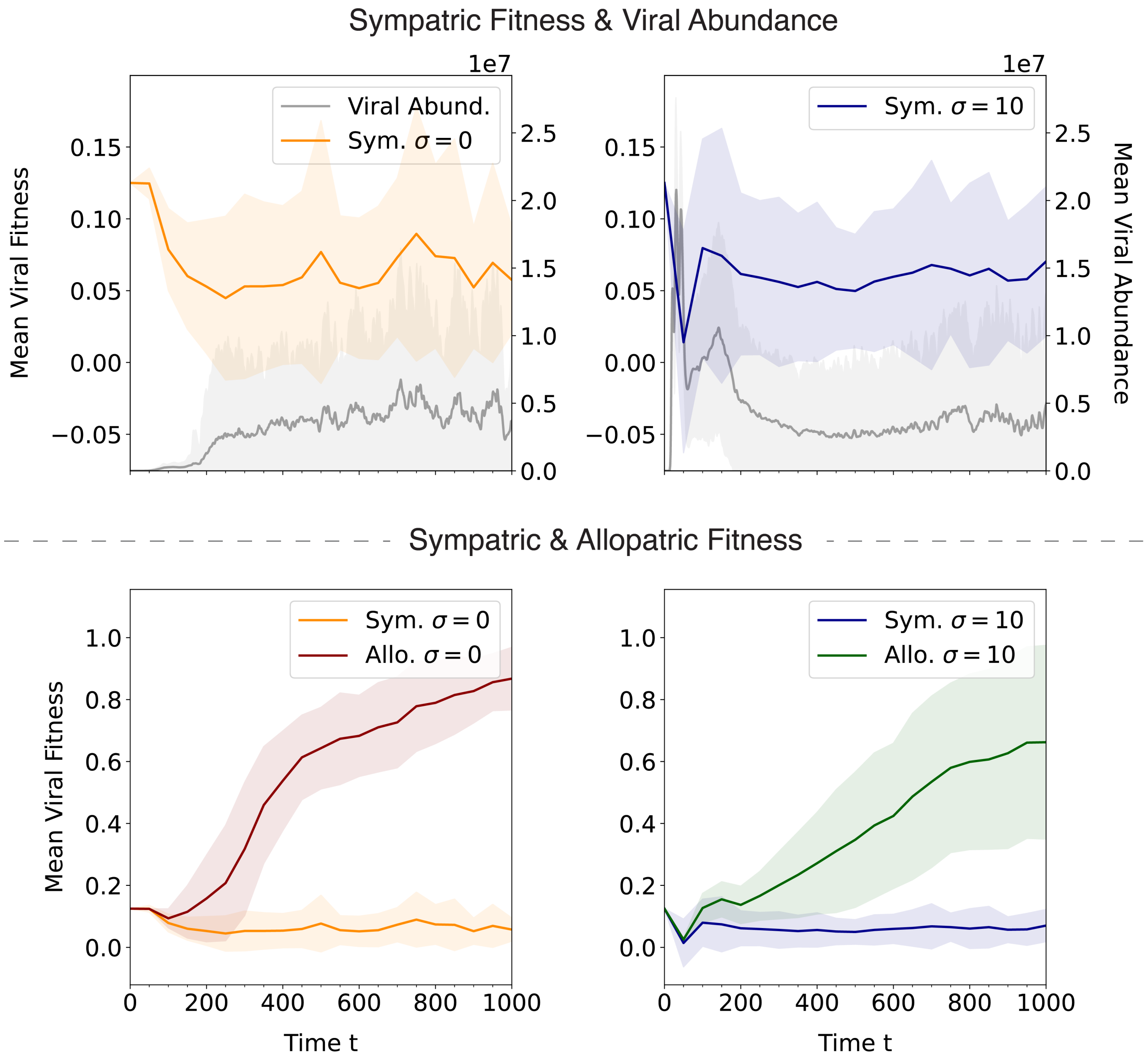


Figure S7: Decomposition of local viral adaptation metric for both extrema of host selection intensities ($\sigma = 0$ and $\sigma = 10$), in treatment I in the absence of competitive-ability mutations. Here, "sympatric" fitness refers to the fitness of the viral population in its locality (replicate), and "allopatric" fitness refers to the fitness of the viral population in the surrounding non-local patches/replicates.
